# Multiple lines of evidence indicate survival of the Ivory-billed Woodpecker in Louisiana

**DOI:** 10.1101/2022.04.06.487399

**Authors:** Steven C. Latta, Mark A. Michaels, Don Scheifler, Thomas C. Michot, Peggy L. Shrum, Patricia Johnson, Jay Tischendorf, Michael Weeks, John Trochet, Bob Ford

## Abstract

The history of decline of the Ivory-billed Woodpecker is long, complex, and controversial. The last widely accepted sighting of this species in continental North America was 1944. Reports of Ivory-billed Woodpeckers have continued, yet in 2021 the U.S. Fish and Wildlife Service proposed declaring the species extinct. We draw on 10 years of search effort, and provide trail camera photos and drone videos suggesting the consistent presence of Ivory-billed Woodpeckers at our study site. Data indicate repeated re-use of foraging sites and core habitat. We offer insights into behaviors of the Ivory-billed Woodpecker that contribute to difficulty in finding this species. We discuss results with regard to the value of accumulated evidence, and what repeated observations may indicate for continued survival of this iconic species.

## Introduction

The history of the decline of the North American population of the Ivory-billed Woodpecker *(Campephilus principalis;* Ivorybill) is long, complex, and controversial (1–3). Currently proposed as extinct (4), the species historically inhabited mature bottomland forests associated with river basins throughout the southeastern United States, with a small, separate population in Cuba (1). Widespread and perhaps very locally common at times, the Ivorybill was severely impacted by collectors, subsistence and other hunters, and cutting of bottomland forests in the U.S. (3, 5). By the late 1930’s, a range-wide search in continental North America resulted in an estimated population of 22 individuals in Florida, South Carolina, and Louisiana (6).

The last widely accepted sighting of an Ivory-billed Woodpecker in North America was in 1944 at the Singer Tract (7), near Tallulah, Louisiana, where Tanner had studied the species (6). Reports of Ivory-billed Woodpeckers continued, however, with authorities estimating as many as 200 sightings since 1944 (8, 9). Many of these reports are from less well-known sources, but some are from game wardens, field biologists, and ornithologists. Observations have also included physical evidence, such as photographs, audio recordings, videos, and a feather (9–13). In 2005, a highly publicized video of a possible Ivory-billed Woodpecker in Arkansas was published (14), but the identification and the survival of the species was strongly debated (15–23). A follow-up, two-year search did not produce additional imagery or documentation widely considered conclusive despite at least 15 reported visual sightings (18). Most recently, evidence suggested that Ivory-billed Woodpeckers were present in the forests along Florida’s Choctawhatchee River (24), and a morphometric analysis of a 2010 photo pointed towards an Ivorybill in Louisiana (25).

These efforts have not resulted in general acceptance of the survival of the species (26). Objections to conclusions of the continued existence of the Ivory-billed Woodpecker among scientists, elements of the birdwatching community, and public media have often focused on two key issues. First, the quality of all reports is said to be so low that they do not offer decisive proof of a living Ivory-billed Woodpecker (16, 27–29). It is argued that a rare bird needs to be documented with a higher standard of evidence and a greater threshold of physical support than routinely adopted for other species; the USFWS defines the objective evidence needed to verify the continued existence of the species as “clear photographs, feathers of demonstrated recent origin, specimens, etc.” (4). The second issue in consideration of the persistence of Ivorybills is the lack of repeatability of observations (28). The assumption is that if a rare resident species is found, then it should be repeatedly re-found, and that if it is not re-found then the original observation or record is inadequate to prove persistence.

Here we draw on 10 years of search effort and provide multiple lines of evidence for the repeated though intermittent presence of Ivory-billed Woodpeckers at our study site in Louisiana.

## Materials and Methods

Our field research took place in bottomland hardwood forests in Louisiana from 2012-2021. Because of the endangered status of the species and ongoing research concerns, we omit specific location details. The search area was defined by perceived habitat quality, previous visual sightings or aural data, and accessibility. The area is a ~93 km^2^ mosaic of bottomlands and uplands. The bottomlands, grown to a variety of hardwoods, occupy a system of drainages and backwaters ~10 km in length, and in breadth from 50 m along some of the smaller feeder streams to ~1.5 km in places along the main stream. This system occurs in a landscape with more remnants of seemingly suitable habitat nearby. The canopy height is ~30 m. Standing and downed dead trees are patchily important components of the landscape. Like almost all bottomland habitat in the southeast, the area has a long history of human use, with most, highly selective lumber extraction having occurred 1890-1940.

Field observations were collected through systematic searching at irregular intervals, with most fieldwork concentrated in the October-May period thought to encompass the breeding season of this species (1). Observational techniques included slowly moving reconnaissance, sitting in place, and stakeouts of key areas, points, or cavities. Boats were not used due to the number and variety of obstructions in the water, reduced mobility, and inability to also handle recording and other equipment. Although skilled, reliable observers reported more than a dozen high quality observations of Ivory-billed Woodpeckers, these are not described here because of the lack of photographic verification. Audio data were also collected through the deployment of ~90 automated recording units. But because of the nature of the known sounds of Ivory-billed Woodpeckers, auditory evidence of the presence of this species is unlikely to be generally persuasive. Likewise, therefore, our audio data are not further described.

Field observations included here were collected through the deployment of trail cameras and the use of drones to record videos. Trail cameras are rugged, weatherproof, programmable, and capable of taking photos automatically at timed intervals or when motion is detected. However, trail cameras are also primarily designed for large animals at ground level and have limited value when shooting upwards to distant treetops where light levels are extreme and motion may not be detected. Nevertheless, trail cameras are one of the few tools available to the Ivorybill searcher. Early images in our search were obtained using Reconyx trail cameras programmed to capture images at 10-sec intervals. Other early images were obtained using Plotwatcher Pro cameras with a dedicated time-lapse device. Plotwatcher software rendered the time lapse in video form; these videos could then be exported in various movie formats, and individual frames exported as jpegs. Image quality was relatively poor, but the review process was simplified. The nature of the time-lapse feature, however, tends to exaggerate the jerkiness of woodpecker movements. More recently, we have used a variety of trail cameras including Moultrie, Bushnell, and Stealth Cam. These were either set on a motion sensitive setting or programmed to take photos at intervals of 5-60 sec.

We placed trail cameras strategically at sites where, a) tight-barked trees appeared to have been scaled, b) trees were damaged or in poor health and expected to die, or c) upright or fallen trees of species favored for feeding by Ivorybills. However, our best results followed placements made when informed by personal visual or aural encounters with Ivorybills. In some instances we focused cameras on the high branches in the mid- to upper-canopy as the strata favored by foraging Ivorybills; however, results showed poor resolution due to distance and light conditions compared to cameras that were focused on lower portions of trunks and fallen branches. Cameras trained to capture images of birds foraging in the mid- to upper-canopy relied on time-lapse programming, while those targeting lower portions of trunks were most often set to a motion sensitive setting.

We sometimes made adjustments to original imagery using standard consumer programs, including Photoshop and Apple Photos. All processing, except cropping, was applied to the entire image; there was no attempt to alter the appearance of individual subject birds. All original still photos presented here were adjusted for contrast and brightness unless otherwise stated.

We have been flying drones equipped with video cameras since July 2019 after published demonstrations of their utility to assess habitat and search for Ivorybills (30), primarily by flying very low over the forest canopy. However, our search method recognizes that Ivorybills regularly fly through the treetops and over the canopy (6), so we hover a drone in place well above the forest, passively filming the treetops to record any birds flying within view of the onboard camera. Hovering the drone at a high altitude, usually just below the Federal Aviation Administration’s maximum height of 400 feet, minimizes disturbance to birds and other wildlife, and creates a relatively stable platform for the camera which results in less blurring of video images than if the drone were moving.

The question of where to aim the camera is an ongoing point of discussion, but our consensus has been that while a directly downward (nadir) view offers the possibility of a very clear view of a passing Ivorybill, it also significantly reduces the field of view and limits the chance of getting a video image of the target bird species. Instead, we film with a shallower (oblique) angle that includes treetops up to 800 m away, increasing the opportunities for an encounter. Finally, where we fly the drones and shoot video is itself informed by many factors, including available habitat, the configuration of habitat on a landscape scale, accessibility of launch sites, permit requirements, and most critically, our history of aural detections and sightings of birds, and locations of foraging signs and cavities.

Flights during 2019 were made with a DJI Mavic 2 Zoom filming with a 4K camera, often using the 2X optical zoom lens. In spring of 2020, we began using Autel Evo II drones with swappable 6K and 8K cameras. Due to a smaller sensor, the 8K camera did not perform well in low light level conditions such as during early morning and on cloudy days, so most videos are now recorded with the 6K camera.

Although the perspective is very different when viewing birds from high above, the process of identifying a species appearing in drone videos is similar to that for birders on the ground, including the basics of assessing the bird’s size and shape, coloration, behavior and habitat. We have been able to distinguish in our footage a wide variety of birds in flight, including Pileated Woodpeckers, and a variety of species of hawks, vultures, ducks, wading birds, and others. We use computer applications such as VLC Media Player, iMovie, Apple Photos, and Topaz Video Enhance AI (for resizing) to analyze our video. We found that analysis was improved significantly with the use of a high quality, high resolution computer screen to view drone videos. As with photo processing, all methods other than cropping were uniformly applied. We also used Photoshop to create a chronophotograph to visualize flights recorded by drones. Chronophotography makes it possible to observe the movement of an object through space in a single still image, thus splitting the perceptual difference between a still image and a moving image, by creating a kind of spatio-temporal snapshot.

## Results and Discussion

We simultaneously deployed 6-19 trail cameras/yr resulting in ~438,000 camera-hrs of activity. Our most important series of trail camera photos followed our sighting of an Ivory-billed Woodpecker landing at ~40 m distance in a live but declining sweetgum *(Liquidambar styraciflua)* tree on 27 October 2019. Trail cameras, nearly continuously deployed on this tree since then, subsequently captured photos of Ivorybills visiting the tree intermittently from at least 29 November 2019 to 10 February 2020, and then again from 12 September 2021 to December 2021.

Trail camera photographs taken on 30 November 2019 and 1 October 2021 at this and a nearby tree, each show a bird with a clear, white saddle on the lower part of the folded wings (Figure 1). Comparative photos of other birds in the same tree (Figure 2), including an unidentified small woodpecker, a Pileated Woodpecker (*Dryocopus pileatus*), and a Red-headed Woodpecker *(Melanerpes erythrocephalus),* confirm the large size of the target bird. While the image quality is too poor for precise measurement, the relatively long neck aspect ratio, proposed as characteristic of the Ivorybill (25), is also highly suggestive. We compare these photographs to one of a known Ivorybill from a separate Cuban population (31) that was also photographed at a considerable distance (Figure 3). The remarkable similarities in the images can be attributed in part to having been shot from ground level, as opposed to almost all existing photos of North American Ivorybills that were obtained from cavity-level blinds (6, 32).

**Fig. 1:**
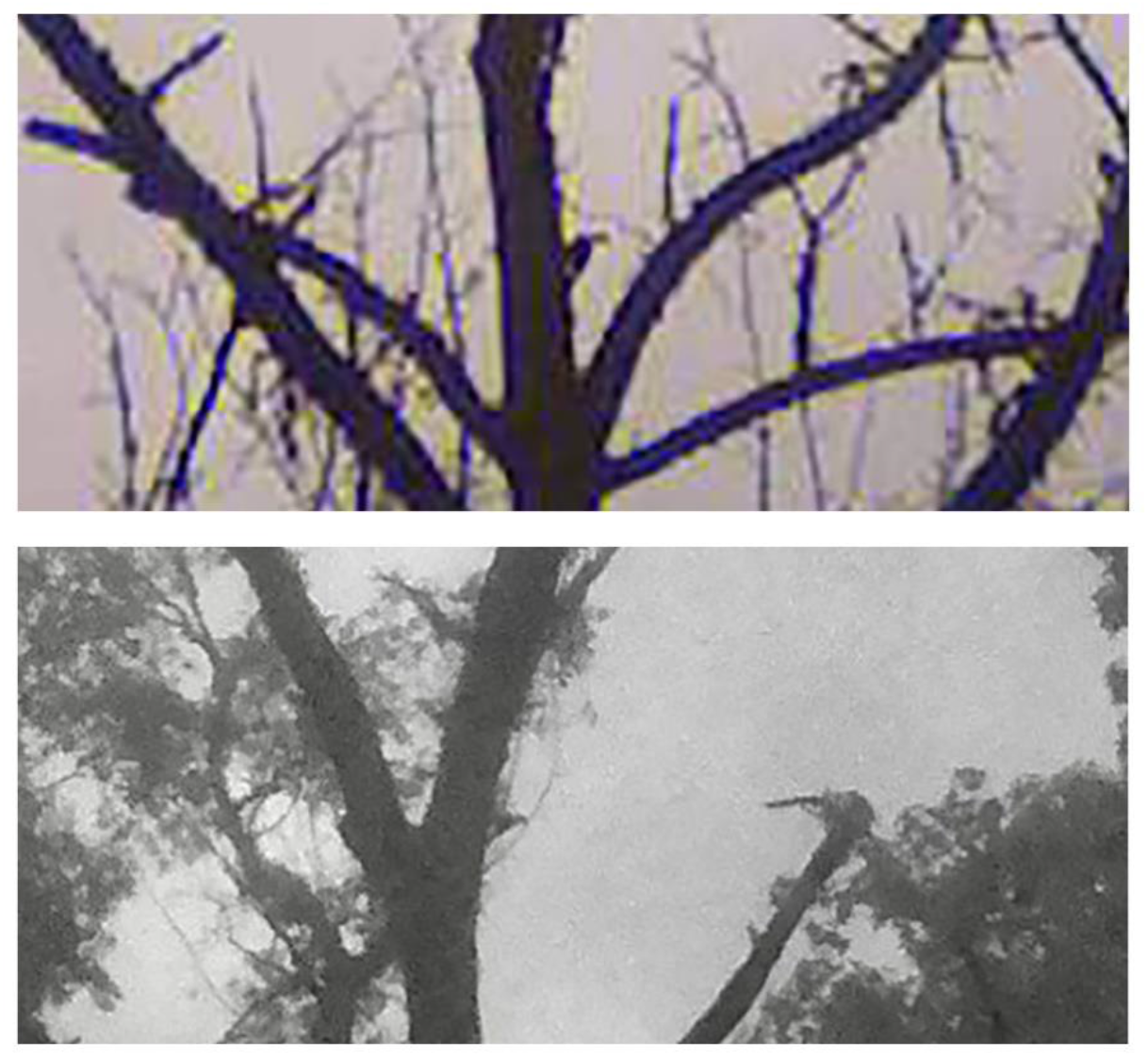
Trail camera photos taken within 50 m of one another on 30 November 2019 (top), and 1 October 2021 (bottom) of apparent Ivory-billed Woodpeckers showing a prominent white saddle present on the lower part of the folded wings. The image from 30 November is extracted from the “video” clip composed of trail cam photographs taken at 5-sec intervals and presented as Supplemental Movie S1.

**Fig. 2:**
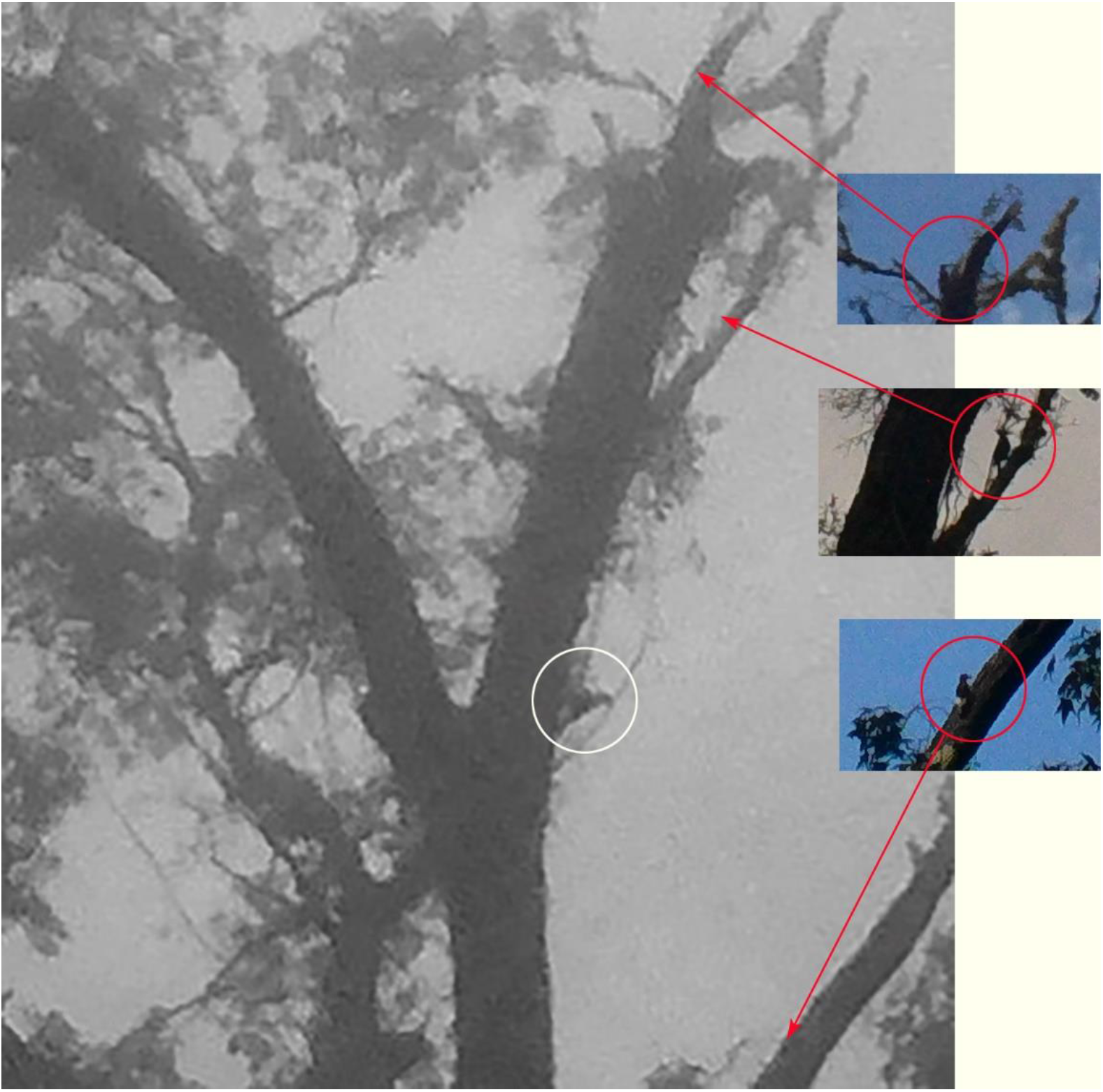
Composite figure comparing the size of three woodpeckers to the apparent Ivory-billed Woodpecker. Inset species were photographed on the same tree, with the same camera in the same place, but at different times. These three images were extracted from their original frames and placed as insets on a fourth frame that shows the Ivorybill on 1 October 2021. All woodpeckers here are depicted at the same scale in their original, unedited size. Arrows point to the location of where each bird was located on the tree. Insets include an unidentified small woodpecker (top), a Pileated Woodpecker (middle), and a Red-headed Woodpecker (bottom). The Ivory-billed Woodpecker is circled in white without an arrow.

**Fig. 3:**
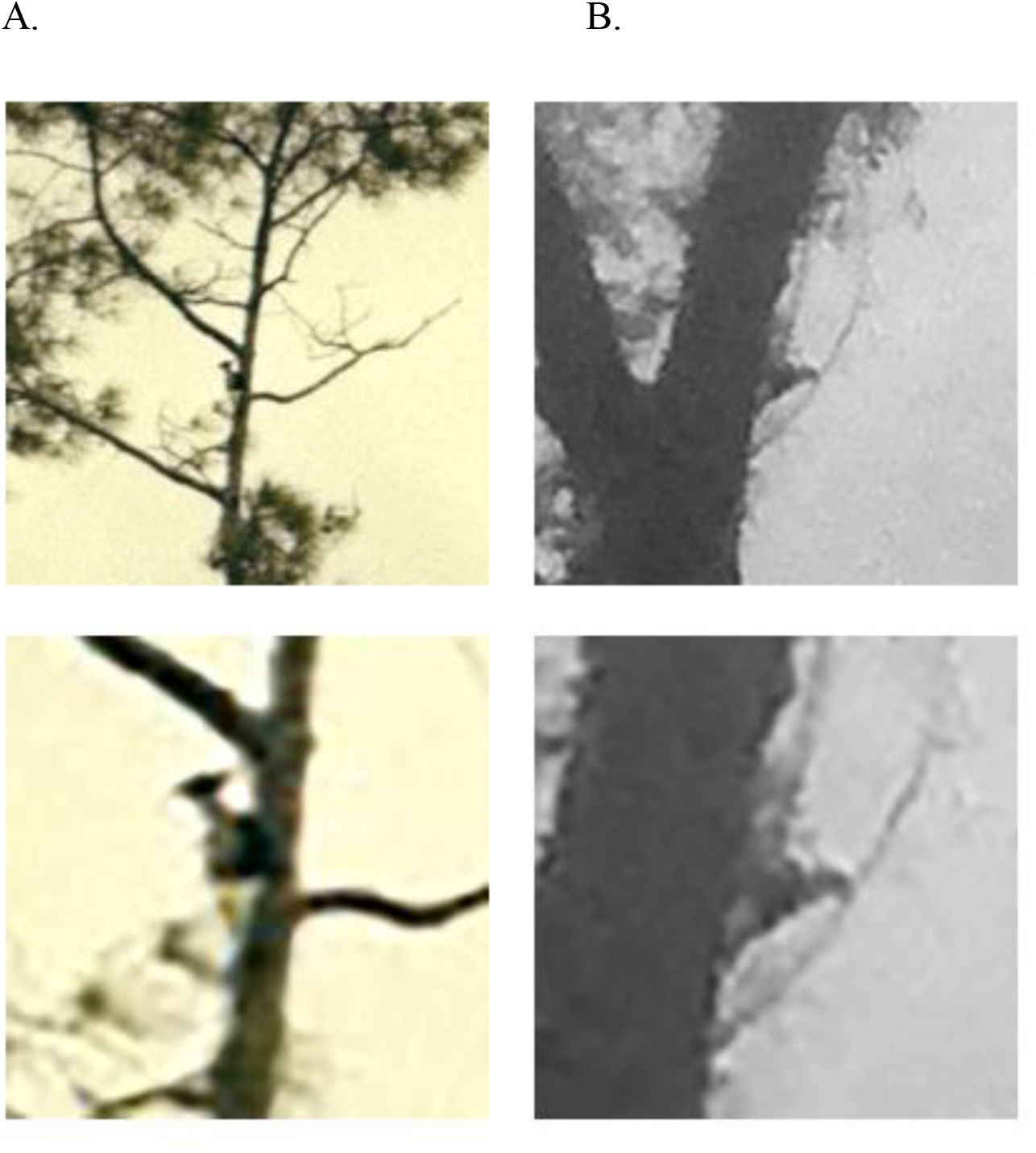
A side-by-side comparison of cropped photos from: A) the unenhanced image of an Ivory-billed Woodpecker from Cuba (31), and B) the original, unretouched Project Principalis photo from Louisiana from 1 October 2021. Each photograph is also shown enlarged and further cropped below each original. These comparisons emphasize the similarities of appearance among Ivorybill images obtained from ground level under challenging field conditions, as opposed to almost all existing photos of North American Ivorybills that were obtained from cavity-level blinds (6, 32).

A series of photographs taken on 12 October 2021 at the same tree occurred in early morning fog (Figure 4). These images of a bird with the typical posture of a large woodpecker are notable for the white saddle. However, the size and prominence of the white saddle changes with the angle of the bird to the camera (Figure 4), as has been previously noted and illustrated (Figure 5) by Jackson (5). We also note that the intersection between the white saddle and the overlying black coverts is uneven in a way that might be expected of feathering. We interpret the image additionally as showing the white dorsal stripe.

**Fig. 4:**
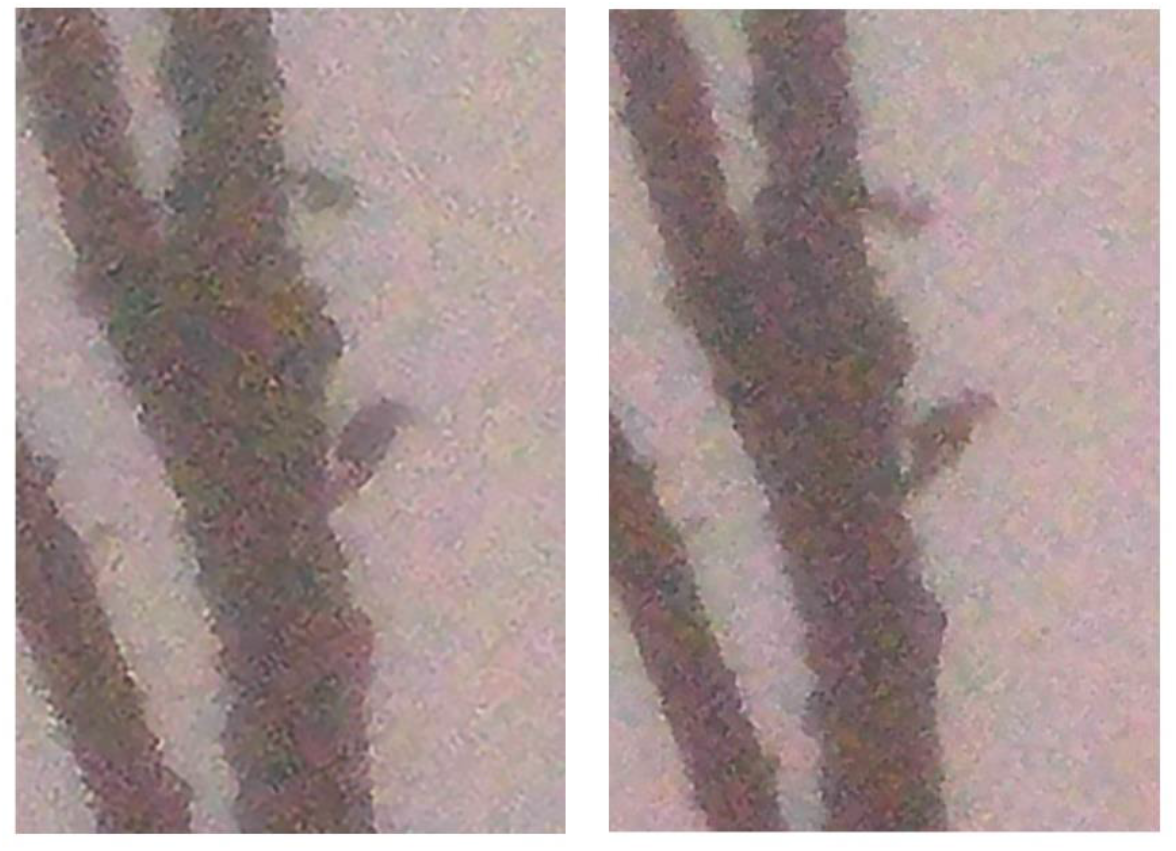
Images of a bird with the typical posture of a large woodpecker, notable for the white saddle despite the heavy, early-morning fog. The size and prominence of the white saddle changes with the angle of the bird to the camera. We also note that an apparent partial white dorsal stripe is present.

**Fig. 5:**
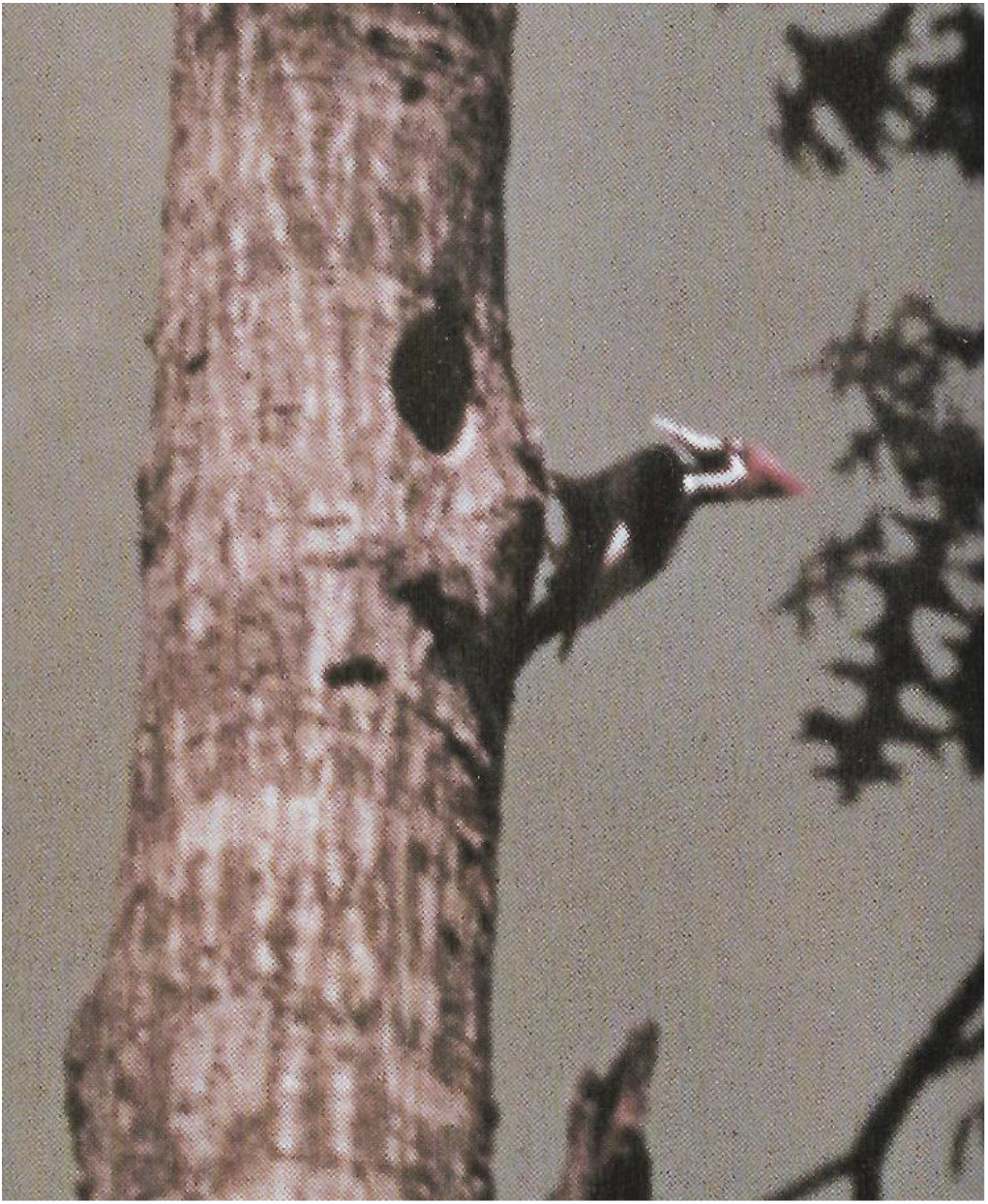
A known Ivory-billed Woodpecker photographed in color in the Singer Tract by James T. Tanner (c. 1939), illustrating how the visibility, size, and prominence of the diagnostic white saddle may be obscured under certain light levels, and depending on the angle of the bird to the camera. Photo by James T. Tanner (5).

Camera images obtained on 14 October 2021 show multiple frames with birds exhibiting distinctive traits of *Campephilus* woodpeckers (Figure 6). A distinctly crested woodpecker with a white saddle, or at least a suggestion of a lighter lower body, is present in many frames. Most intriguing is that these images depict the distinctive morphological adaptations of the feet and legs of *Campephilus* woodpeckers as compared with *Dryocopus* woodpeckers like the Pileated Woodpecker (33). The phenotypically similar Pileated is one of the most unspecialized of the truly arboreal woodpeckers, while the *Campephilus* woodpeckers are characterized by pamprodactyly, a pedal morphology that enables the forward rotation of all four toes (33). The specialized modifications in the highly arboreal Ivory-billed Woodpecker are not so much in the structure of the toes as in the position of the legs. The feet are held outward from the body and are directed diagonally upward and sidewise (Figure 7), with both feet wide apart and more anterior relative to the body (33, 34). Usually the angle between the tarsi and the horizontal plane is ≤45°, and often seem to be pressed against the tree trunk. This is very different from the condition seen in most woodpeckers, as, for example, the Pileated Woodpecker, where the legs are held more or less beneath the pelvic girdle, the joints are fully flexed, and the tarsi are held well away from the tree trunk. This generally results also in a more obtuse angle of the intertarsal joint (where the leg bends between the tibiotarsus and the tarsometatarsus), and is evidence of the better scansorial adaptations of the Ivory-billed Woodpecker compared to the Pileated Woodpecker (33). This obtuse angle is visible from a distance, is readily seen in our images (Figures 6, 7), and can be a useful identification clue in situations where lighting or distance makes it hard to observe plumage details with clarity (35). Combined with feet extended diagonally upward and to the side of the body, the stance of the birds appearing here are consistent with that of a *Campephilus* sp.

**Fig. 6:**
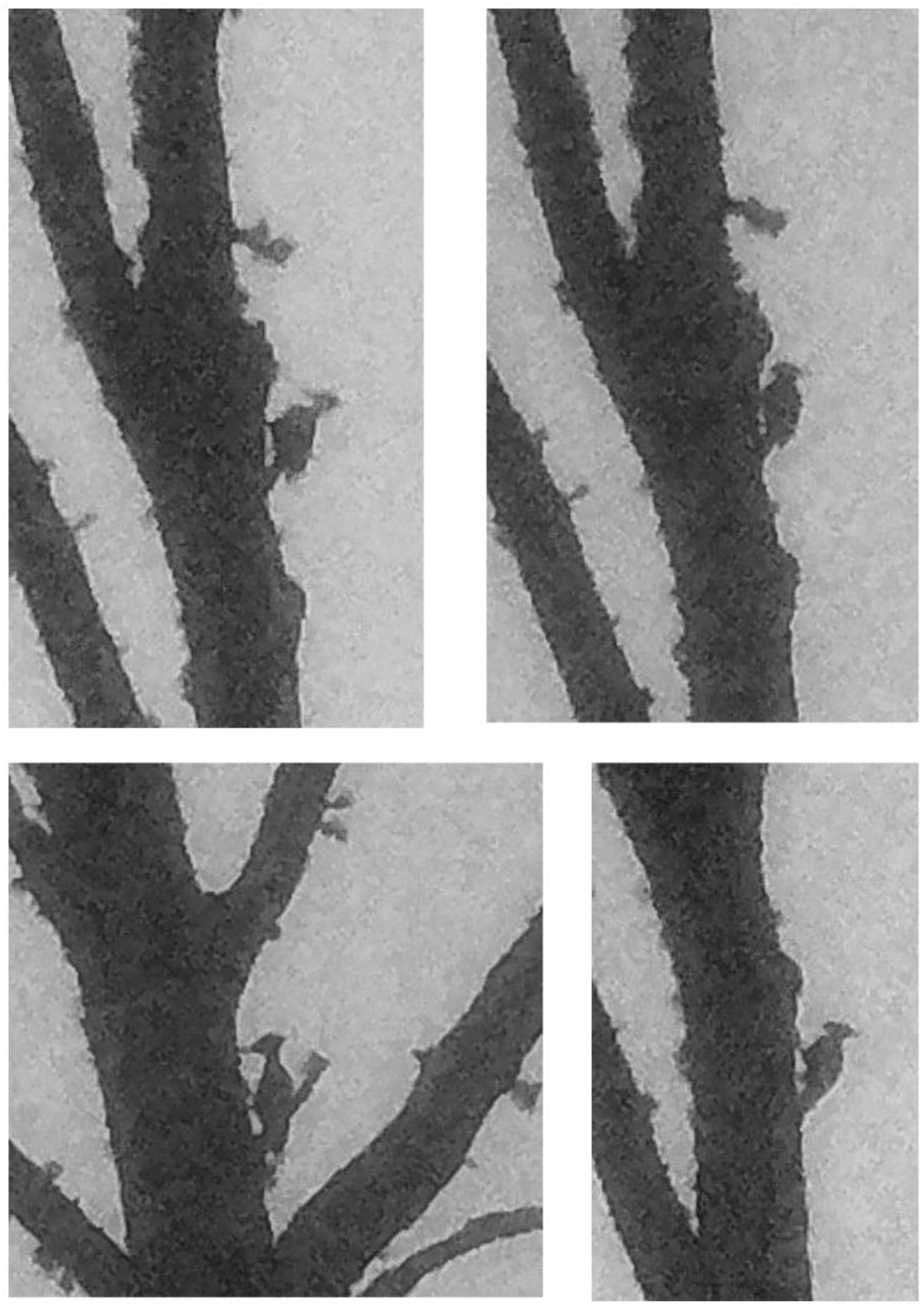
Project Principalis images from 14 October 2021 illustrating the specialized modifications in the position of the legs in the highly arboreal Ivory-billed Woodpecker. The feet are held to the side of the body and are directed diagonally upward and sidewise, with both feet wide apart and forward. Usually the angle between the tarsi and the horizontal plane is ≤45° and there is an obtuse angle of the intertarsal joint. While a white saddle is not obvious in these early morning, very misty silhouettes, several images suggest its presence.

**Fig. 7:**
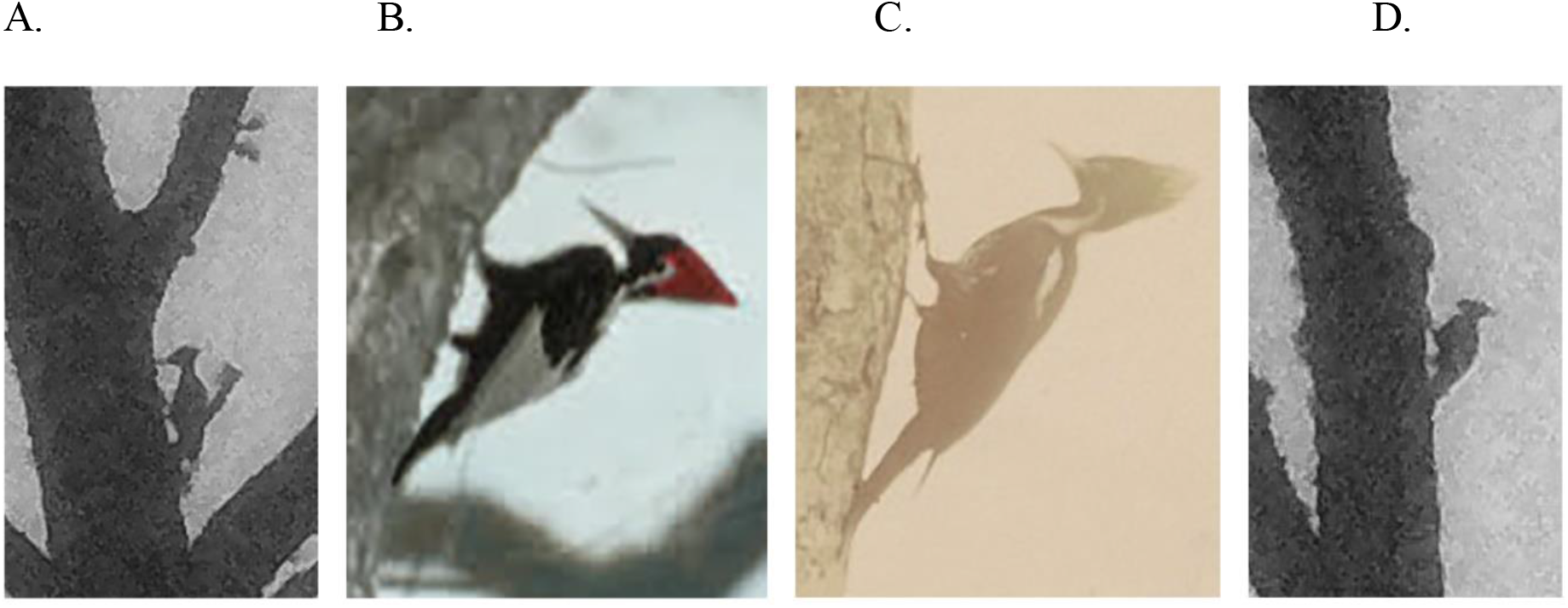
Comparison of photographs taken of apparent Ivory-billed Woodpeckers in Louisiana from this study (A, D), with a colorized Ivory-billed Woodpecker, also from Louisiana, but taken by Arthur A. Allen in 1935 (B), and a Pale-billed Woodpecker *(Campephilus guatemalensis)* taken in Central America (C), also from the Allen Collection. Birds in all photos share the characteristic posture imposed by the unique structure of the *Campephilus* leg and feet. Feet are held to the side of the body and are directed diagonally upward and sidewise, with both feet wide apart and forward. The angle between the tarsi and the horizontal plane is ≤45° and there is an obtuse angle of the intertarsal joint. Photos (B) and (C) are from the James T. Tanner, and the Arthur A. Allen papers, respectively, courtesy Division of Rare and Manuscript Collections, Cornell University Library.

A final set of trail camera images offers further behavioral clues to the identification of these birds. On 9 January 2020, an apparent male-female pair of Ivory-billed Woodpeckers was photographed (Figure 8). This image shows one bird with an apparent red crest, another with an apparent black crest, and a prominent white saddle on the male. One of the photo sequences we find most compelling, however, was obtained on 30 November 2019. These trail camera photos involve what appears to be a foraging family group. When viewed in succession (Supplemental Movie S1), the resulting “video” clip appears to show three large woodpeckers moving and foraging together. The “video” is composed of individual trail camera photographs taken automatically every 5 sec. Although distance and lighting are difficult, a white saddle can be clearly seen in multiple frames, including a frame extracted and reproduced in Figure 1 showing a woodpecker with a prominent white saddle on the lower part of the folded wings. We note also the proximity of the three birds to one another in the “video”, and their foraging behavior, including movements throughout the tree: on the bole and major branches, and even on smaller branches. Foraging appears to be very active and even acrobatic at times, with birds clinging to the tops, sides, and undersides of the branches. We recorded very similar foraging behavior on the same tree on 12 October 2021, with very active and acrobatic movements across the tree, including smaller branches (Supplemental Movie S2). The apparent ~2-yr gap in foraging by Ivorybills on this nearly continuously monitored tree is interesting, suggesting that there is an intermittency in woodpecker movements, or more likely, in the phenology of beetle prey and their larvae. Continued, long-term monitoring of trees utilized by Ivorybills is warranted to better understand foraging patterns.

**Fig. 8:**
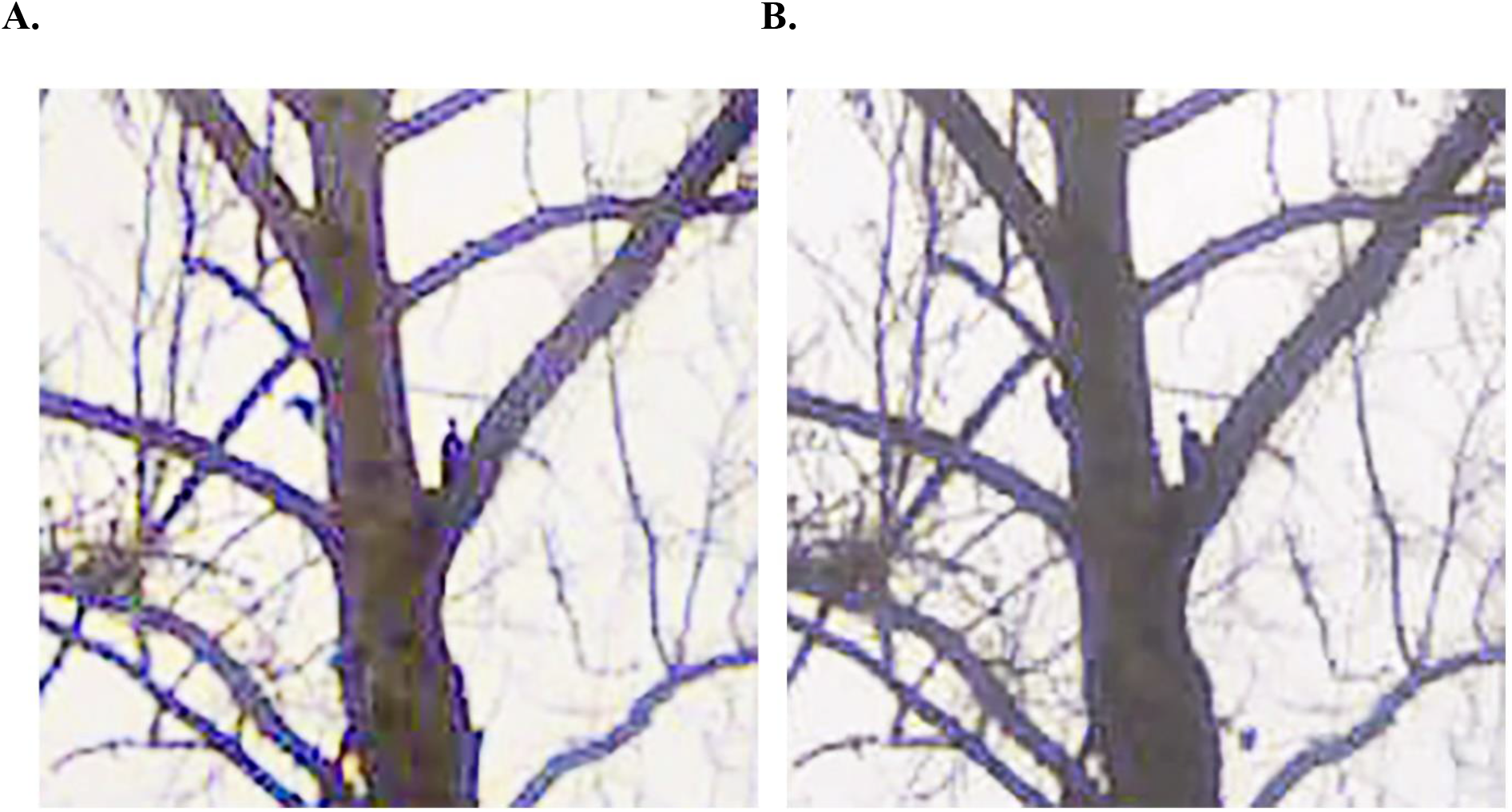
An apparent male-female pair of Ivory-billed Woodpeckers photographed on 9 January 2020 with the bird on the left showing an apparent red crest, and the bird on the right showing an apparent black crest. Among North American woodpeckers, only the female Ivorybill has a black crest. A prominent saddle is present in Figure 8A on the bird on the left, and a partial white dorsal stripe is present in Figure 8A on the bird on the right.

Intraspecific behavior may also support the identification of these birds as Ivory-billed Woodpeckers. Ivory-bills reportedly show no indication of being strongly territorial (6). In the Singer Tract, home ranges did not appear to be defended during the breeding season, and wandering birds that were encountered seemed to be tolerated by resident birds. In addition, Sonny Boy, the male offspring that Tanner banded in 1937, remained with his family group for a full two years after fledging (32). By contrast, the Pileated Woodpecker generally appears to be territorial year-round, only tolerating birds from other territories at distances of >100 m (36). Adult Pileateds typically drive young away from the territory in the fall, often as early as September, but anecdotal reports do exist of three Pileateds together during winter months (36). Our observations of three birds appearing just a few meters apart, well after a presumed fledging period, and for an extended time, is more consistent with an Ivorybill family group than an unusual Pileated or mixed-species group, but should not be considered definitive. However, considering that we see white on the wings of birds in successive frames (Figures 1), even at considerable distance and under poor lighting conditions, we are confident that these sequences include Ivory-billed Woodpeckers.

In addition to the evidence of a family group, the observed foraging behavior is distinctly unlike that of a Pileated Woodpecker. Pileateds select large diameter trees (36, 37), and dead trees are used out of proportion to availability (37). Large rectangular excavations are characteristic; these can be >30 cm in length (36). Although Pileateds may also glean and peck, their bark scaling behavior is a distinctly uncommon activity in Louisiana bottomlands (37). Pileated Woodpecker foraging tactics are rather slow and methodical, and concentrated on the bole and major branches of large trees, as the species avoids trees in smaller size classes (37). The foraging style of the Ivory-billed Woodpecker seems to be largely undescribed, other than the importance of scaling of bark of hardwoods (6). It is unclear from the literature whether foraging as active as we document is typical of the species, but our subsequent careful inspection of the smaller branches of the tree where the Ivorybills were photographed did reveal extensive scaling of even the smaller branches in the canopy. Furthermore, photographs taken by Tanner in 1939 similarly reveal a group of three Ivorybills foraging together on a tree at the same time, while also documenting that the three birds were also less than a meter apart from one another (32).

We also used drones to document the presence of Ivory-billed Woodpeckers at our study site. We made ~2,590 drone flights and recorded ~864 hr of video from July 2019 – October 2021. We recorded several drone videos in February 2020 that captured images of large birds with black on the leading edge of the wings and white on the trailing edges. These videos were taken in an area where we had had recent sightings and had recorded vocalizations suggestive of Ivorybills. In a video that was recorded on 25 February 2020, an apparent Ivory-billed Woodpecker crosses the field of view and banks upward and to the right before landing on a large, emergent tree. To illustrate the approach and landing, we present the landing sequence as a chronophotograph (Figure 9). The black leading edge with white trailing edge is repeated in multiple frames. While the white in several images looks more like a tail, similar to that of a Bald Eagle (*Haliaeetus leucocephalus*), other images make it clear that the white is on the back of the wings. In addition, a Bald Eagle with a white tail would always show a bold white head, which never appears on this bird. We believe that the false appearance of a white tail in some frames is a result of the positioning of the wings closer together toward the posterior of the bird during the return wing stroke. The bird also appears to land in a distinctly woodpecker-like fashion on a vertical tree trunk or slightly inclined limb with a quick upward swoop and a few braking wing-beats (6). The bird’s white saddle then appears to be visible on the trunk of the tree after landing. Based on shade patterns on other trees, the bird is not in full sun at the time of landing so the apparent white saddle is not an aberration caused by reflected direct sunlight.

**Fig. 9:**
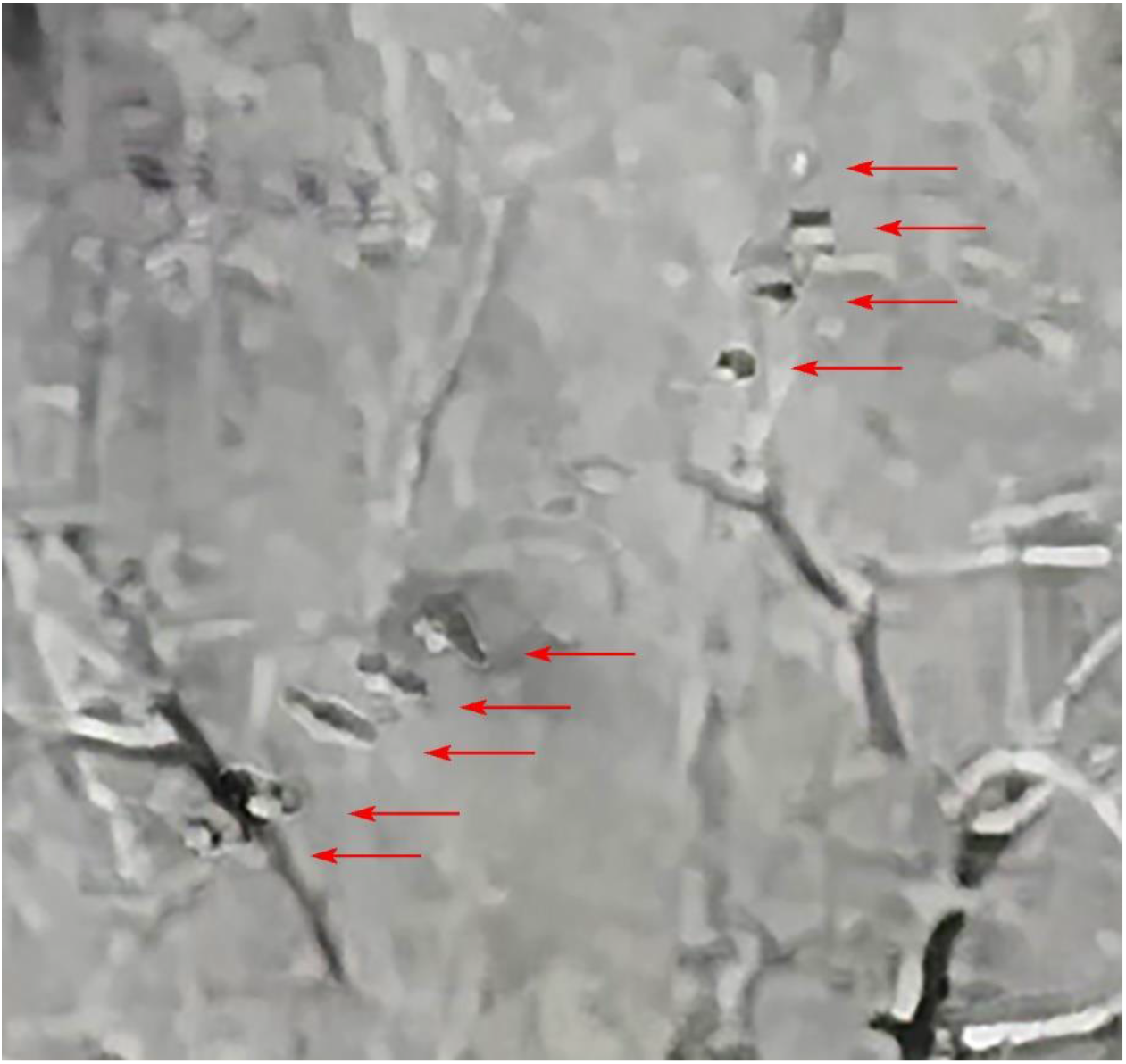
A chronophotograph of a landing sequence showing an apparent Ivory-billed Woodpecker in February 2020. The arrows point to individual images of the bird. The direction of flight is generally from lower left to upper right across the field of view.

A second informative video from 23 December 2019 shows two large birds in flight above the treetops. Shortly after entering the view of our camera, the first bird appears to engage in flight bounding, a behavior in which a bird stops flapping by temporarily folding its wings onto its back. For a moment, it speeds missile-like before flapping again. As the bird alternates between flapping and flight bounding, the result is a flight path that appears slightly undulating, which is common among woodpeckers. Although the historical literature about the Ivory-billed Woodpecker does not mention flight bounding, a 1939 photograph of an adult Ivorybill flying overhead (Figure 10) is evidence that there are moments when the wings are folded on top of the body. An additional clue to what an Ivory-billed Woodpecker’s bounding flight should look like from a drone can be found in a 1956 video (38) of the closely-related Imperial Woodpecker in Mexico (Figure 11). The video offers a partial side view showing some of the white patch on the bird’s back during flight bounding.

**Fig. 10:**
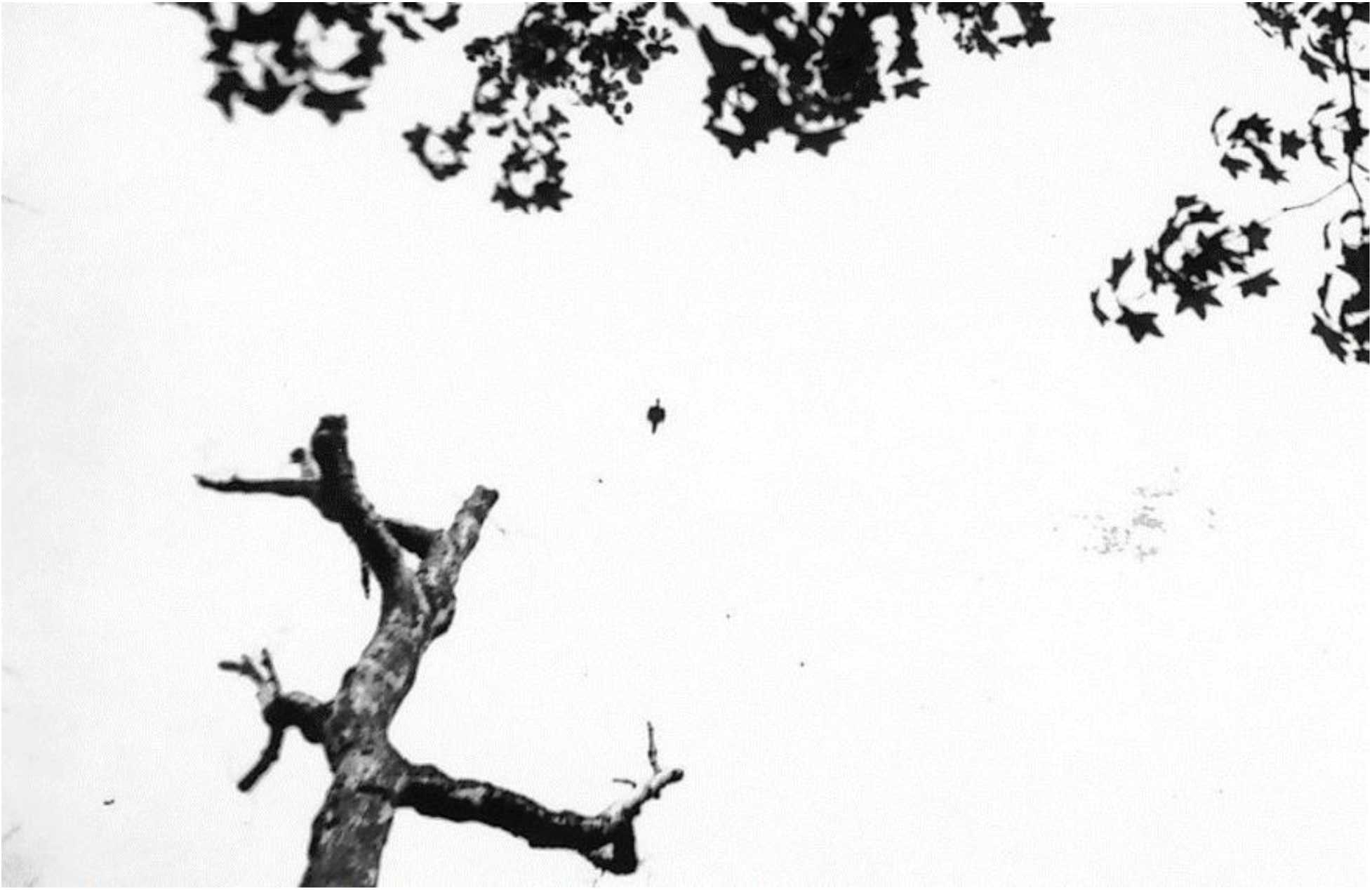
A photograph by James T. Tanner from April 1939 of an Ivory-billed Woodpecker demonstrating flight bounding by this species. Photograph courtesy LSU Digital Libraries, Louisiana, and the Lower Mississippi Valley Collections, Louisiana State University, Baton Rouge, Louisiana.

**Fig. 11:**
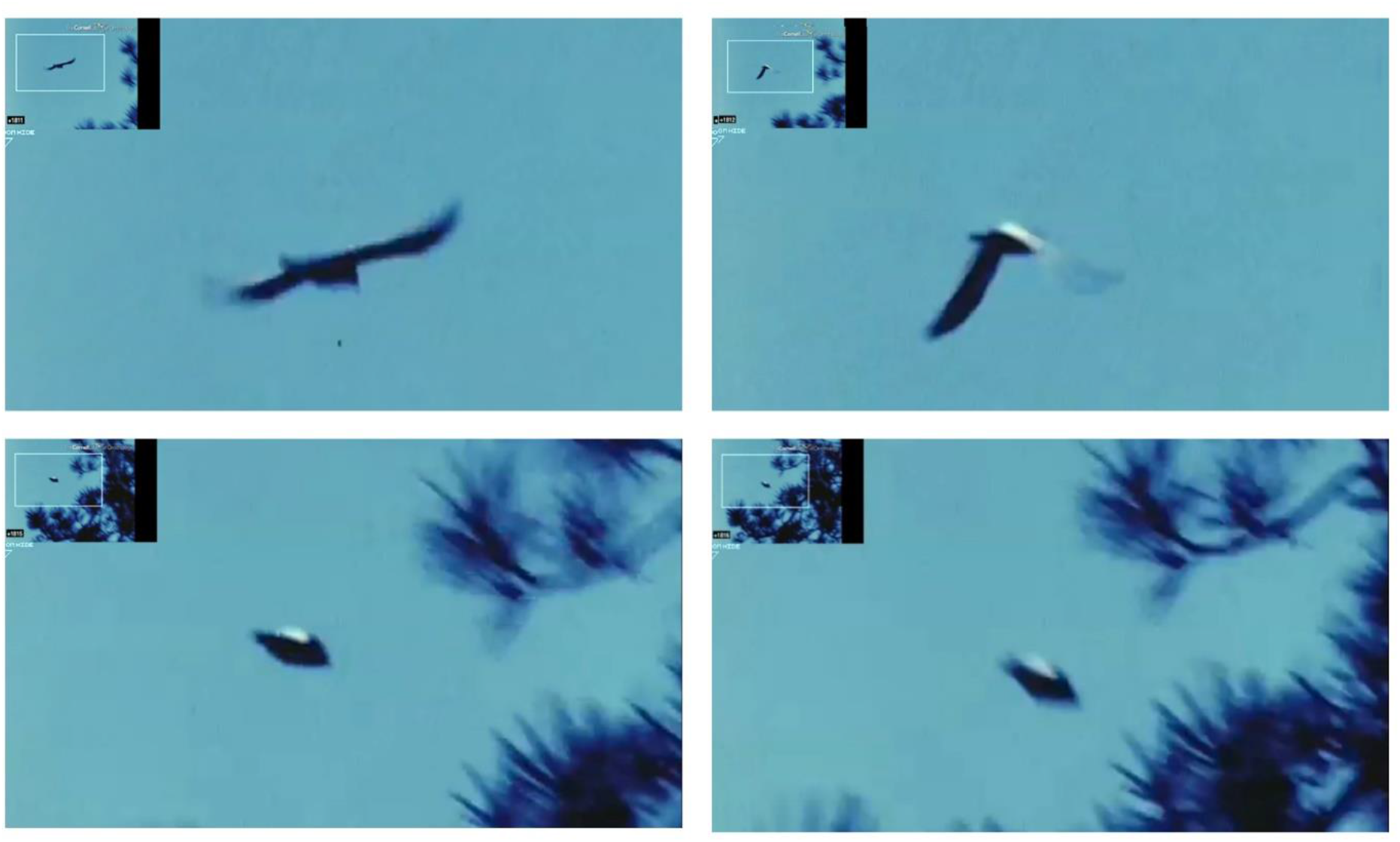
Stills (left to right, top to bottom) from a video shot from a Mexican hillside by William L. Rhein of the closely related Imperial Woodpecker *(Campephilus imperialis),* demonstrating flight bounding by this species (38). Video available from Macaulay Library at the Cornell Lab, *Campephilus imperialis* (ML#61027).

Our December 2019 clip similarly shows a large dark-colored bird as it flies away from the camera and to the right; our video perspective is from above and behind the subject bird. In the frames reproduced in Figure 12, the white plumage appears along the trailing edge of the wing as the bird begins to fold its wings inward. In subsequent frames, the white condenses into a bright patch as the wings complete their fold onto the bird’s back, then appearing very nearly as a typical white saddle of an Ivory-billed Woodpecker.

**Fig. 12.**
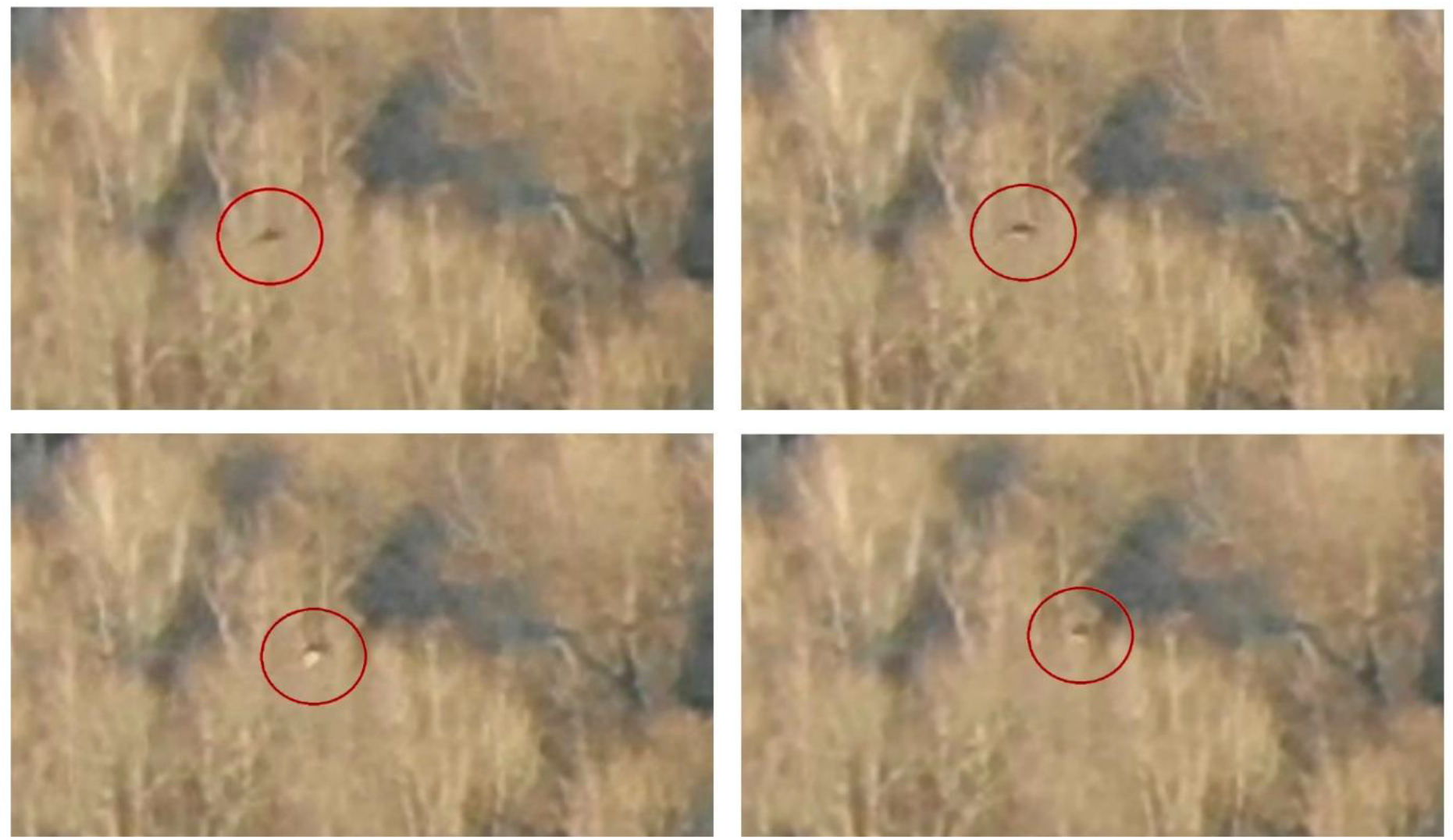
Stills (left to right, top to bottom) from a drone video shot in Louisiana of an apparent Ivory-billed Woodpecker demonstrating flight bounding by this species. White plumage appears along the trailing edge of the wing. In subsequent frames, as the bird begins to fold its wings inward, the white condenses into a bright patch. The white then appears very nearly as a typical white saddle as the wings complete their fold onto the bird’s back.

Two compelling behavioral features are derived from the drone videos. One is the height at which the bird appears to travel. Flights above the treetops extending “for a half mile or more” and “dodging the trees with very little deviation from their course” have been described for the Ivorybill in North America (6); Ivorybills have also been reported to fly high over canyons and treetops in Cuba (39). A second compelling feature is the high speed and direct flight of Ivorybills. Direct flight was previously noted in the Ivorybill (6, 40), while Pileated Woodpecker flight is characterized as “rather slow, but vigorous and direct” (36).

The data presented here offer no doubt that the multiple images and videos are those of a large woodpecker. Our opinion that these images represent Ivory-billed Woodpeckers is based on the appearance of broad white saddles, white trailing edges on the wings of birds in flight, a white stripe along the side of the neck, a heavy, light-colored bill, a unique morphology of the legs and feet, and a pair of birds with one suggesting a black crest. With the number of images available, some with multiple individuals present, one can also safely eliminate unusual aberrations and leucistic oddities that are sometimes posited as explanations for a large, woodpecker-like bird with extensive white plumage. Related to aberrations are defects or distortion of the video, frequently recognized as foreign artifacts. However, artifacts are far less of an issue with recent HD video technology, and should be of much less significance in evaluating video shot in 4K or 6K HD as we have presented here.

We note that through many trail camera encounters at these distances, many photographs remain ambiguous. Some frames clearly show the white saddle of the Ivorybill, and these field marks can be seen in successive or multiple frames. However, in some cases, successive frames may show no white visible for the same birds. As mentioned above, we suggest that lighting conditions and position of the bird accounts for the absence of white in these cases, as documented by some historic photographs of known Ivorybills in the Singer Tract (Figure 5), where the white saddle is almost totally obscured. In addition, the angle of the camera to the bird affects the amount of white appearing in a photograph. In this case, we are shooting an apparent Ivory-billed Woodpecker from underneath and at considerable distance when the trail camera is near ground level and the canopy is ~30 m high. These conditions contrast markedly with those which produced almost all photographs in the historical record; all but a handful of existing photographs of known Ivory-billed Woodpeckers were taken from a blind placed at nearly eyelevel to a nesting cavity in the Singer Tract. These ideal photographic conditions have, we believe, impacted expectations of what photographs are likely under field conditions.

The variety of evidence we have gathered over many years indicates repeated re-use of foraging sites and core habitat, and offers unusual repeatability of detections of the species. Lack of repeatability of observations has been raised in the past to dismiss purported Ivorybill sightings. For example, countering claims around the Luneau video from Arkansas, critics suggested that, “experience with other rare birds, especially resident species, suggests that any valid sighting should very quickly lead to more sightings” (28). This criticism was lodged despite the fact that the Luneau video followed a series of sightings, and was itself followed by additional sightings and acoustic recordings (14). Repeatability in our observations is seen at a variety of scales. All of the observations reported here took place in a single forested block and a single watershed. Almost all of the encounters reported here occurred within 1.6 km of one another; and the majority of the best trail camera photos were taken over two, 3-month periods on the same tree.

We believe that our observations contribute to a clearer understanding of the twin problems of why the Ivory-billed Woodpecker has been so difficult to detect and to relocate over the past 80 years. These issues begin with the misperception that, if present, the Ivorybill is relatively easy to find – a misperception that extends at least as far back as Tanner (6). Tanner was a meticulous observer, but he apparently never located an Ivory-billed Woodpecker outside the Singer Tract, despite his numerous searches throughout the southeast (6, 41). Tanner noted that, “The difficulty of finding the birds, even when their whereabouts was known … limited the number of observations (6).” Nonetheless, the misperception emerged, sometimes fueled by Tanner himself, that the Ivorybill should be noisy and easy to find. But this was entirely based on a single family group during the nesting season, and in later years Tanner acknowledged that this was usually not, in fact, the case (42).

Misperceptions on the ease of finding the Ivorybill extend to the frequent argument that in the modern era it is unlikely that a large, distinctive woodpecker could escape the sights, cameras, and recorders, of birdwatchers and other people who are recreating or working outdoors in remote areas (43–45). For example, even with the popularity of birdwatching, birdwatchers are not everywhere. The eBird citizen science program (https://ebird.org/home) has amassed >33 million checklists (46). While the most thorough coverage occurs in North America, modeling of the range and relative abundance of individual species at a 3 km spatial resolution resulted in areas of “no predictions” because there was an insufficient number of qualifying checklists to assess whether a species was present or absent (47). While eBird checklists occur at easily accessible places in the vicinity of our study area, no eBird checklists occur from within our specific area.

The authenticity of reports from non-scientists, hunters, fishermen, and rural residents, who may be the most likely people to access habitats such as those occupied by the Ivorybill, are often dismissed. Though often keen and knowledgeable observers of their natural world, their observations of rare or unusual species are frequently devalued relative to the science-based perspectives of researchers (48, 49). Increasingly, however, new approaches to science methodology recognize that local people often have a very intimate relation with the environment and natural resources. The closeness of these relations and dependencies is such that these people have a very particular and detailed knowledge of local environmental conditions and ecological relations that is of value to science (50).

Beyond the questions of detection and documentation, our data offer insights into how the ecology and behavior of the Ivory-billed Woodpecker would contribute to the difficulty in finding or re-finding this species. We know that the Ivorybill inhabits some of the most difficult to access habitat in the U.S., and that mature bottomland forests are a core component of that habitat. Behaviorally, our observations showing long, high-altitude flights by single birds and pairs are suggestive of a species with a vast home range, and accustomed to utilizing dispersed and likely fragmented habitats. Home ranges may vary seasonally, but the Ivorybill pair studied in the Singer Tract may have had a range up to four miles or more in diameter (6). Ivorybills have also been reported to wander over even greater distances and to cross cutover and otherwise unsuitable habitat (6, 51). Large home ranges, distant wandering across unsuitable habitat, and high flights all suggest the need, and the willingness, of the Ivorybill to travel widely for patchily distributed resources. That may be to search for a suitable roost site, but more frequently, it may be related to exploitation of abundant but ephemeral food sources that occur in dead or dying trees or branches.

## Conclusion

The habitat as described above applies to many other places in the American Southeast. The continuing survival of Ivory-billed Woodpeckers in Louisiana has conservation management implications not only in that state, but also widely within the historic range of the species. We expect that Ivorybills persist in some of these other places also, if not permanently then episodically. Their numbers cannot be expected to improve unless many more large and continuous bottomland hardwood forests are actively or passively managed to exhibit old growth characteristics. Forested tracts must be large enough so that ecological changes caused by natural catastrophic events such as fires, floods, or hurricanes, as at Congaree National Park owing to Hurricane Hugo in 1989, will allow surviving Ivory-billed Woodpeckers opportunity for a diversity of habitats. Only then, can there be an expectation of a larger number of populations or subpopulations of this iconic species.

The report contained here is not the end of our efforts. We are encouraged and energized by what we have accomplished. We are optimistic that technologies will continue to improve our outcomes, including documentation through environmental DNA and other physical evidence. We believe that our intentional and systematic survey design is paying off through complementary lines of investigation. Our findings begin to tell a larger story not just of whether the Ivory-billed woodpecker persists in Louisiana, but how it has survived and why its survival has been so difficult to document. Finally, we also believe that our methodologies can be translated to other sites, thus offering opportunities for additional documentation of the species. Our findings, and the inferences drawn from them, suggest an increasingly hopeful future for the Ivory-billed Woodpecker.

## Acknowledgements

We dedicate this paper to the memory of Frank Wiley, co-founder of Project Coyote, predecessor of Project Principalis. Also to the late Bob Russell, John Trochet’s amiable and dedicated search partner for more than a decade in Louisiana, Arkansas, South Carolina, and Florida. We thank the numerous volunteers who assisted (and continue to aid) in this search, and we thank the landowners who provided permissions to access the sites. We thankfully acknowledge the participation and field contributions of Project Principalis associates Tom Foti, Erik Hendrickson, Richard Martin, Geoffrey McMullen, Stephan Pagans, and Philip Vanbergen. Peter Shefler helped create figures. The images appearing in this publication are copyrighted and owned by their individual owners. All rights reserved. We thank the Macaulay Library at the Cornell Laboratory of Ornithology for receipt of media; the Division of Rare and Manuscript Collections, Cornell University Library, for use of photographs from the James T. Tanner papers, and the Arthur A. Allen papers; and the LSU Digital Libraries, Louisiana and Lower Mississippi Valley Collections, Louisiana State University Libraries, Baton Rouge, Louisiana, for photos. The findings and conclusions in this article are those of the author(s) and do not necessarily represent the views of the U.S. Fish and Wildlife Service.

## Supporting Movies

**Movie S1:** “Video” clip composed of trail cam photographs taken at 5-sec intervals showing three large woodpeckers foraging together on 30 November 2019. Unlike adult Pileated Woodpeckers that are territorial year-round and typically drive young away from the territory as early as September, Ivory-billed Woodpeckers reportedly show no indication of being strongly territorial, and offspring have remained with family groups for a full two years after fledging. Figure 1, showing birds with apronounced white saddles, is extracted from this “video” clip.

**Movie S2:** Time lapse “video” clip composed of trail cam photographs taken at 30-sec intervals showing two large woodpeckers foraging together on 12 October 2021. The images are in silhouette, and field marks are not visible until one of the birds is higher on the tree, at which point the white saddle on the back can be seen. As in Movie S1, these large, crested woodpeckers demonstrate very active foraging movements on a branch at lower left before moving up the tree. Foraging is acrobatic; the birds hang from vines and cling to the tops, sides, and undersides of the branches.

